# Mesenchymal stem cells protect retinal ganglion cells from degeneration via mitochondrial donation

**DOI:** 10.1101/393959

**Authors:** Dan Jiang, Hong Feng, Zhao Zhang, Bin Yan, Ling Chen, Chuiyan Ma, Cheng Li, Shuo Han, Yuelin Zhang, Peikai Chen, Hung-Fat Tse, Qingling Fu, Kin Chiu, Qizhou Lian

**Author notes:** **Corresponding authors:** Dr. Qizhou Lian, Dr. Kin Chiu, Dr.Qingling Fu.

## Abstract

Retinal ganglion cell (RGC) degeneration is extremely hard to repair or regenerate and is often coupled with mitochondrial dysfunction. Mesenchymal stem cells (MSCs)-based treatment has been demonstrated beneficial for RGC against degeneration. However, underlying mechanisms of MSC-provided RGC protection are largely unknown other than neuropectective paracrine actions. In this study, we sought to investigate whether mitochondrial donation can preserve RGC functions, in a mitochondrial *Ndufs4* deficient mouse model of RGC degeneration. The results revealed intravitreal transplanted by induced pluripotent stem cell derived-MSCs (iPSC-MSC) could donate their mitochondria through crossing inner limited membrane to host RGCs. Furthermore, the donated mitochondria effectively protected against RGC death and largely preserved retinal function in *Ndufs4*-KO mice. Importantly, the protective effects of mitochondrial donation from MSCs were associated with management of pro-inflammatory cytokines. Our data identified a novel role of MSCs-mitochondrial donation in protection of RGC from degeneration, and highlight a viable therapeutic strategy by manipulating stem cell mitochondrial donation for the treatment of retina degeneration in future.

## Introduction

Exogenous administration of mesenchymal stem cells (MSC) is protective in retinal degeneration, including glaucoma and Leber’s hereditary optic neuropathy (LHON) (Inoue et al., 2007; Johnson et al., 2010a; Ng et al., 2014; Yu-Wai-Man et al., 2014), suggesting that MSCs might be neuroprotective in this context. Several on-going clinical trials indicate the feasibility and safety of using MSCs for the treatment of neuromyelitis optical disorders (www.clinicaltrials.gov, NCT01920867; NCT02249676) and age-related macular degeneration (NCT02016508) although the underlying mechanisms of any benefit remain unclear. MSCs might improve retinal cell survival via cell replacement, although there is little convincing evidence that MSCs are capable of functional neural differentiation (Joe and Gregory-Evans, 2010; Kicic et al., 2003; Phinney and Prockop, 2007; Ratajczak et al., 2004; Tomita et al., 2006). Conventionally, MSC transplantation has long been thought to provide neuroprotective paracrine factors for retinal protection (Johnson et al., 2010a; Manuguerra-Gagne et al., 2013; Mead et al., 2015; Torrente and Polli, 2008). Nonetheless, use of paracrine factor-based therapies has often produced inconsistent and even conflicting results in treatment studies (Inoue et al., 2007; Li et al., 2009). Thus, the paracrine effects of MSCs do not fully explain the mechanisms of MSC-modulated retinal protection.

MSCs are capable of transferring their mitochondria to pheochromocytoma PC12 cells, lung airway cells, corneal cells and cardiomyocytes, and improve mitochondrial bioenergetics (Islam et al., 2012; Jiang et al., 2016; Plotnikov et al., 2008; Rustom et al., 2004; Spees et al., 2006). Recently it has been reported that MSCs can transfer mitochondria to astrocytes and restore the bioenergetics of the recipient cells and stimulate their proliferation (Babenko et al., 2018). We considered the alternative possibility that the protective effect of MSCs involves mitochondrial transfer to neural cells in the retina such as retinal ganglion cells (RGC). RGCs are responsible for propagating signals derived from visual stimuli in the photoreceptor cells in the eye to the lateral geniculate body/superior colliculus in the brain, through their long axons (London et al., 2013; Rodieck, 1965; Zhang et al., 2016). Due to the natural complexity of RGCs in neuronal development, axon guidance, and complex synaptic connections, they are almost impossible to repair or replace once damaged (Aguayo et al., 1990; Almasieh et al., 2012; Goldberg et al., 2002; Hilla et al., 2017; Levkovitch-Verbin et al., 2001). Another challenge is that it is thought unlikely that intravitreal injected cells can pass through the inner limiting membrane of the retina (Quigley and Iglesia, 2004). We have previously reported that compared with adult bone marrow-derived mesenchymal stem cells, induced pluripotent stem cell-derived mesenchymal stem cells (iPSC-MSCs) can more efficiently donate functional mitochondria to rejuvenate injured heart, lung and corneal tissues (Jiang et al., 2016; Li et al., 2018; Zhang et al., 2016). Hence, it is of great import to investigate whether iPSC-MSCs can transfer mitochondria to RGCs and whether such a transfer is therapeutic.

We studied these issues using a *Ndufs4* knockout (KO) mouse model with defined mitochondrial complex I deficiency (Yu et al., 2015). We reported an unexpected acute effect of intravitreal iPSC-MSCs transplantation on the rescue of RGCs in this model through direct mitochondrial transfer. Importantly, the protective effects of mitochondrial donation from MSCs were associated with management of pro-inflammatory cytokines. Our findings challenge the current belief that protection of RGCs by MSCs is merely through their paracrine effect and suggest that the functional mitochondrion of MSCs can pass through the inner limiting membrane and halt mitochondrial damage-provoked RGC death, enabling us to better understand stem cell therapy in retinal mitochondrial disorders and to develop a novel strategy for retinal repair by manipulating intercellular mitochondrial transfer.

## Results

### Retinal ganglion cell degeneration in Ndufs4 deficient mice

To evaluate the onset and development of RGC damage and so determine the optimal period for therapeutic intervention, we first evaluated the changes to RGCs over time in Ndufs4 KO mice. Given a life span of up to 9 weeks for KO mice (Irwin et al., 2013), retinal pathological changes were examined between week 3 and week 7. To determine the loss of RGCs, counting cells that were Brn3a positive, an RGC marker, was performed on mounted whole retinas of KO and WT mice. The number of Brn3a positive RGCs in KO mice retinas declined significantly at week 6 and week 7 when compared to WT (Figure 1A). No significant changes were observed in ERG when *Ndufs4* KO mice were compared with WT mice at week 3 (Figure 1B). Nonetheless compared with WT mice, significant decreases in b-wave and PhNR amplitude were observed at week 4 in KO mice (p<0.05 WT vs. KO). Morphometric analysis of the retinal section revealed that the thickness of the inner plexiform layer (IPL) of both the central and peripheral retina had declined significantly in KO mice by postnatal week 7 (Figure 1C). In addition, since high levels of glial fibrillary acidic protein (GFAP) has been associated with Müller cell activation accompanies with RGC loss (Kurihara et al., 2006), we examined GFAP expression at different times in KO mice. At week 3, GFAP positive cells were observed only in Müller end-feet and astrocytes in the nerve fiber layer of retinas in both WT and KO mice (Figure EV1). At week 4, the expression of GFAP in retinas of KO mice reached into the inner plexiform layer (IPL). By week 7, an increasing fluorescent signal of GFAP was observed to axial extent throughout the retina of KO mice (Figure EV1).

**Figure 1.**
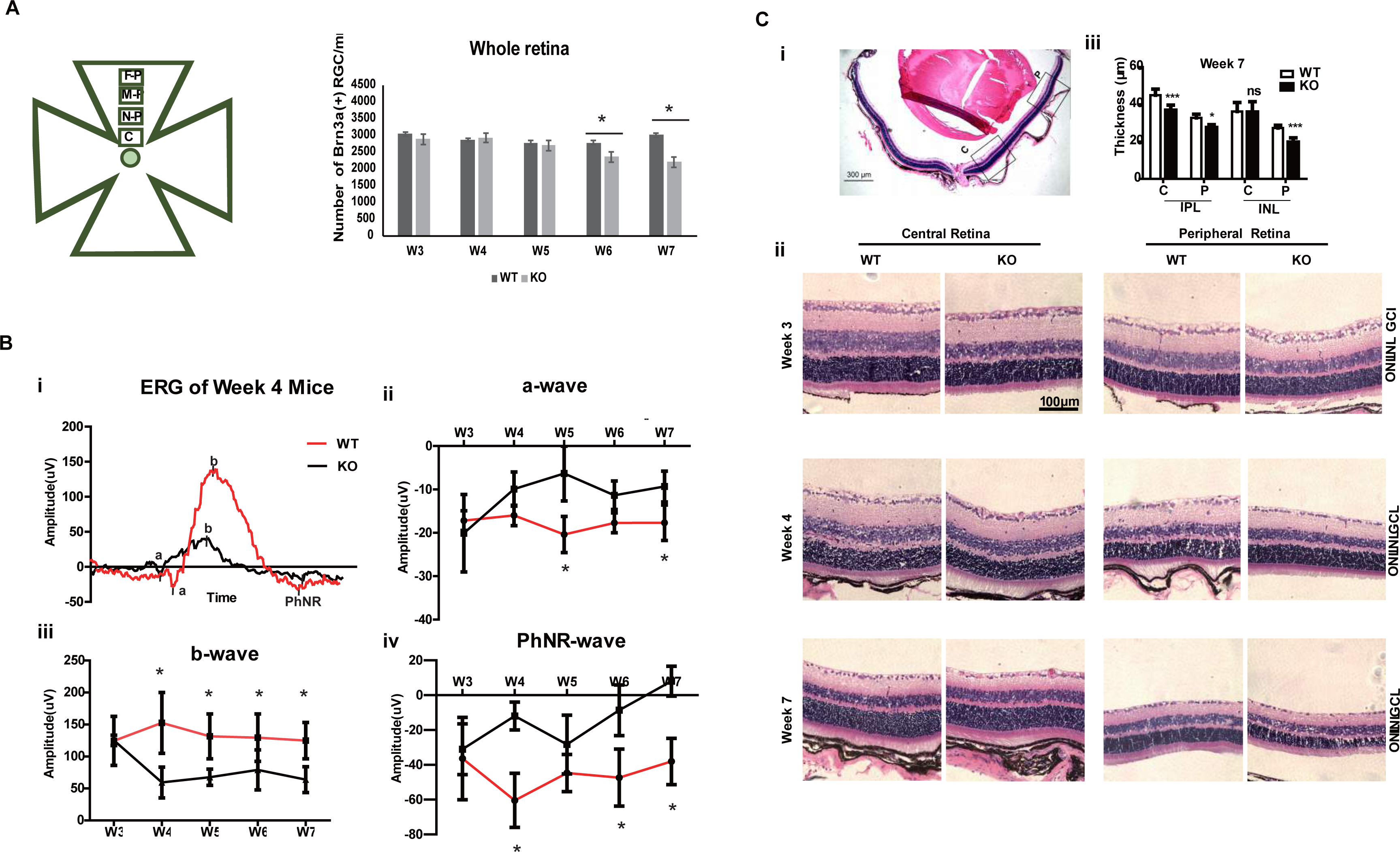
Dissection of Retinal Ganglion Cells Degeneration in Ndufs4 deficient Mice. **A** Analysis of Brn3a (+) RGCs in whole mounted retina. Schematic diagram of a whole flat mounted retina shows four analyzed regions: central retinal (C), near-peripheral (N-P), mid-peripheral (M-P) and far-peripheral (F-P) retina. The average number of Brn3a positive cells in KO and WT mice was counted from W3 to W7. (n□=□5 eyes/group). **B** Electroretinogram of Ndufs4 KO and WT mice. Representative traces recorded from a single animal demonstrating the a-wave, b-wave and PhNR components at week 4. Mean amplitude of a-wave, b-wave and PhNR from week 3 to 7 at the flash intensity (16.60 scd/m2) (n=10). **C** Hematoxylin and Rosin (H&E) analysis of retinal sections from Ndufs4 mice. i) Images were taken from two areas of the retina: central retina (C) and peripheral retina (P). ii) Representative H&E images of central and peripheral retina of 3, 4, and 7 week old KO and WT mice. iii shows the thickness of INL as well as IPL of central and peripheral retina of KO and WT mice’ retina at week 7 (n=5). (inner nuclear layer (INL), plexiform layer thinning (IPL), outer nuclear layer (ONL), retinal ganglion cell layer (GCL)). Data information: In (A–C), data are presented as mean ± SD. *p≤0.5, **p≤0.01, ***p≤0.001, ns, not significant (Student’s t-test).

Taken together, a time course-based ERG analysis revealed dysfunction of retinal ganglion cells started at week 4, RGC density decrease in the flat-mounted retina was detectable at week 6, and significant reduced retinal thickness in morphometric analysis at week 7. Abnormal activation of Müller cells was detectable by week 4 and extensively distributed throughout the whole retina by week 7. These data support the hypothesis that mitochondrial complex I dysfunction in the retina results in RGC functional loss and death, similar to the phenotype of Leber’s hereditary Optic Neuropathy. On or before week 3 at the very early phase of RGC damage may be a treatment window to prevent RGC loss and functional degeneration. For this reason, we performed intravitreal injection at week 3 and examined mice 96 hours, 1 week and 4 weeks post iPSC-MSC transplantation (Figure EV2).

### Detection of mitochondrial transfer from iPSC-MSCs to retina

To determine whether intravitreal delivery of iPSC-MSCs could transfer mitochondria to retinal cells and pass through the inner limiting membrane *in vivo,* mitochondria of iPSC-MSCs were genetically labeled with LV-Mito-GFP prior to intravitreal injection(Zhang et al., 2016). In the whole mount retina, GFP positive mitochondria from iPSC-MSCs were detected within TUBB3 positive mouse RGCs (Figure 2A). Z-stake reconstructed images and video of the retina (Figure 2B-C and Movie EV1) clearly showed that most mitochondria of iPSC-MSCs were translocated within the ganglion cell layer (GCL). In addition, only a few mitochondria were detected within the inner nuclear layer (INL) and inner plexiform layer (IPL) (indicated by triangular arrowheads) (Figure 2B-C). This indicates that cells in the GCL directly took up mitochondria from iPSC-MSCs. We also detected the transferred mitochondria in primary isolated RGCs during co-culture with MSCs *ex vivo* (Figure EV3).

**Figure 2.**
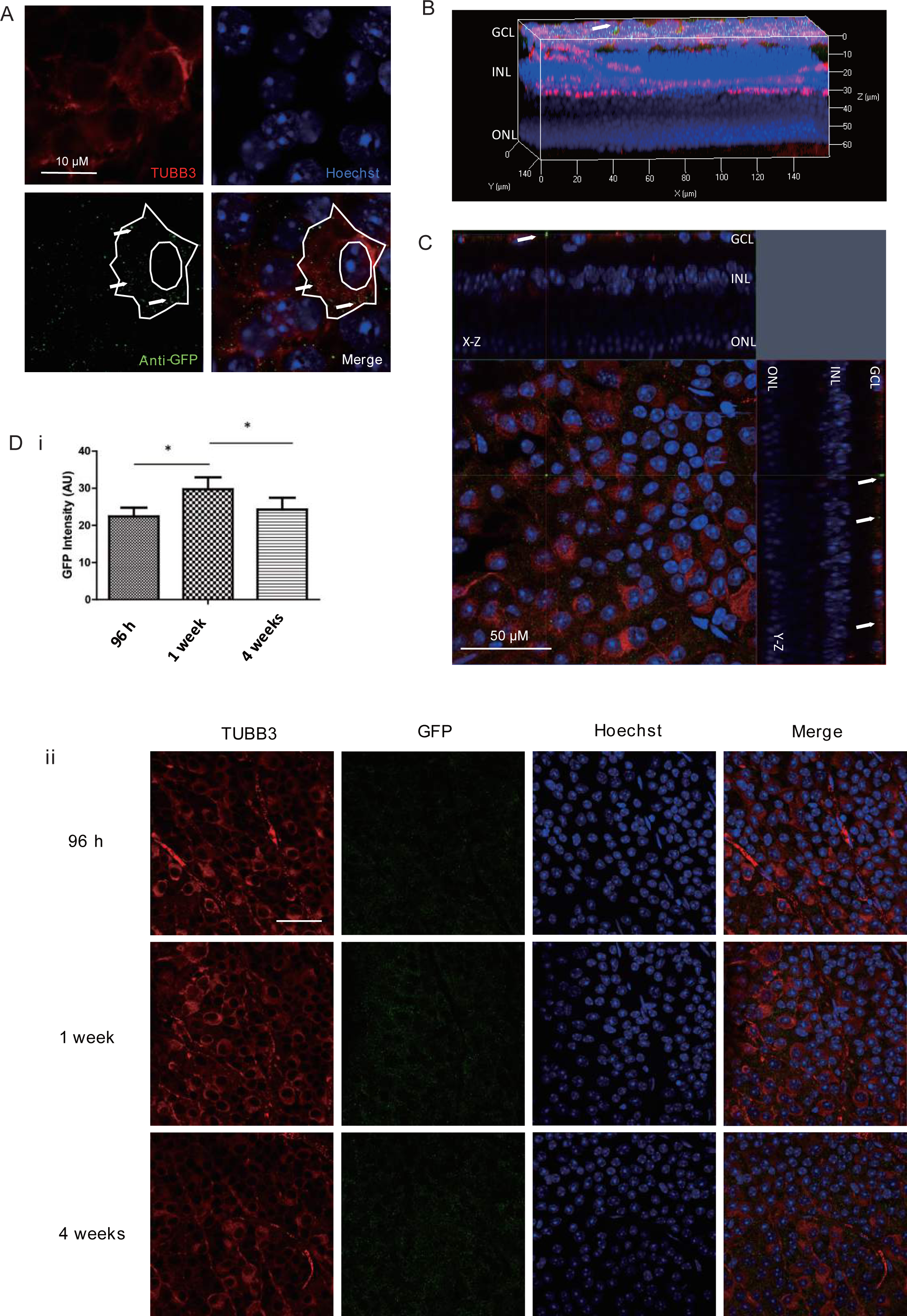
Detection of mitochondria transfer from MSCs to retina in RGC. **A** At one-week post iPSC-MSCs administration, immunofluorescence was performed in whole mount retina to evaluate mitochondrial transfer of iPSC-MSCs by staining with anti-GFP (Green), RGC marker TUBB3 (Red) and Hoechst (Blue). GFP positive mitochondria could be detected within the RGC. A white line delineates a cellular and nucleus outline and arrows show the GFP positive mitochondria within the RGC. **B** Z-stack reconstructed images show the location of the mitochondria from iPSC-MSCs in retina (arrow). **C** XY-axial image of ganglion cell layer (GCL) shows GFP positive mitochondria within RGC layer (indicated by arrows). XZ-axial and YZ-axial images show the different sublayers. Scale bars: 50 μM. **D** i) On 96 h, 1 week and 4 weeks post iPSC-MSC administration, GFP intensity was analyzed. (n=10, Error bar represent SD, p-values calculated using One-way ANOVA, * p≤0.05). ii) immunofluorescence image show GFP in whole mount retina. Scale bars: 40 μM

To examine transfer efficiency of mitochondria, mitochondria within retina were analyzed at 96 h, 1 week and 4 weeks post iPSC-MSC administration by immunostaining assay against anti-GFP (488), TUBB3(594), and Hoechst (405) staining. Mitochondrial transfer from iPSC-MSCs to the retina was evident as early as 96 hours after treatment and GFP intensity was significantly increased 1 week post iPSC-MSC administration (Figure 2D). By 4 weeks, no MSC left in the vitreous chamber on top of the inner limiting membrane. However, GFP positive mitochondria were still detectable within the RGL (Figure 2D). To exclude non-specificity of immunostaining against GFP, another set of antibodies was used including Brn3a (488), anti-human mitochondria (anti-Mito, 594) and DAPI (405) (Figure EV4). Similarly, human mitochondria were detected and were most abundant 1-week post iPSC-MSC administration, compared with 96 hours and 4 weeks (Figure 2D & Figure EV4).

### Penetration of mitochondria but not MSCs into retina

To determine whether injected iPSC-MSCs could penetrate the inner limiting membrane and migrate to the RGC layer or just transfer healthy mitochondria into retinal cells, mice retinas were isolated from the underlying RPE and snap frozen for PCR. Vitreous body was carefully separated from retina. Four pairs of primers designed for human specific nuclear DNA(H-nDNA), human specific mitochondrial DNA (H-mDNA), mice specific nuclear DNA(M-nDNA) and specific mitochondrial DNA (M-mDNA) were used to detect DNA information within the retinal tissue (Zhang et al., 2016). In all the iPSC-MSC injected mice eyes, human mitochondrial DNA was detected within mice retinas but no human nuclear DNA was detected at any of the analyzed timepoints (Figure 3A). To exclude the possibility of contamination of human cells in mice retina in PCR assay, we performed immunostaining against human-nuclear antibody (HNA) to detect human-specific antibody in retinal sections. We found positive HNA (Human nuclear marker) only in the vitreous cavity but not any layers of the retina (Figure 3B-C). These results indicate that mitochondrial transfer from iPSC-MSCs to the retina occurred as early as 96hrs following iPSC-MSC injection and remained within the retina 4 weeks after administration. Of note, after intravitreal injection, only mitochondria of iPSC-MSCs could be transferred into the mice retina while no iPSC-MSCs could pass through inner limiting member and integrated into the retina of mice.

**Figure 3.**
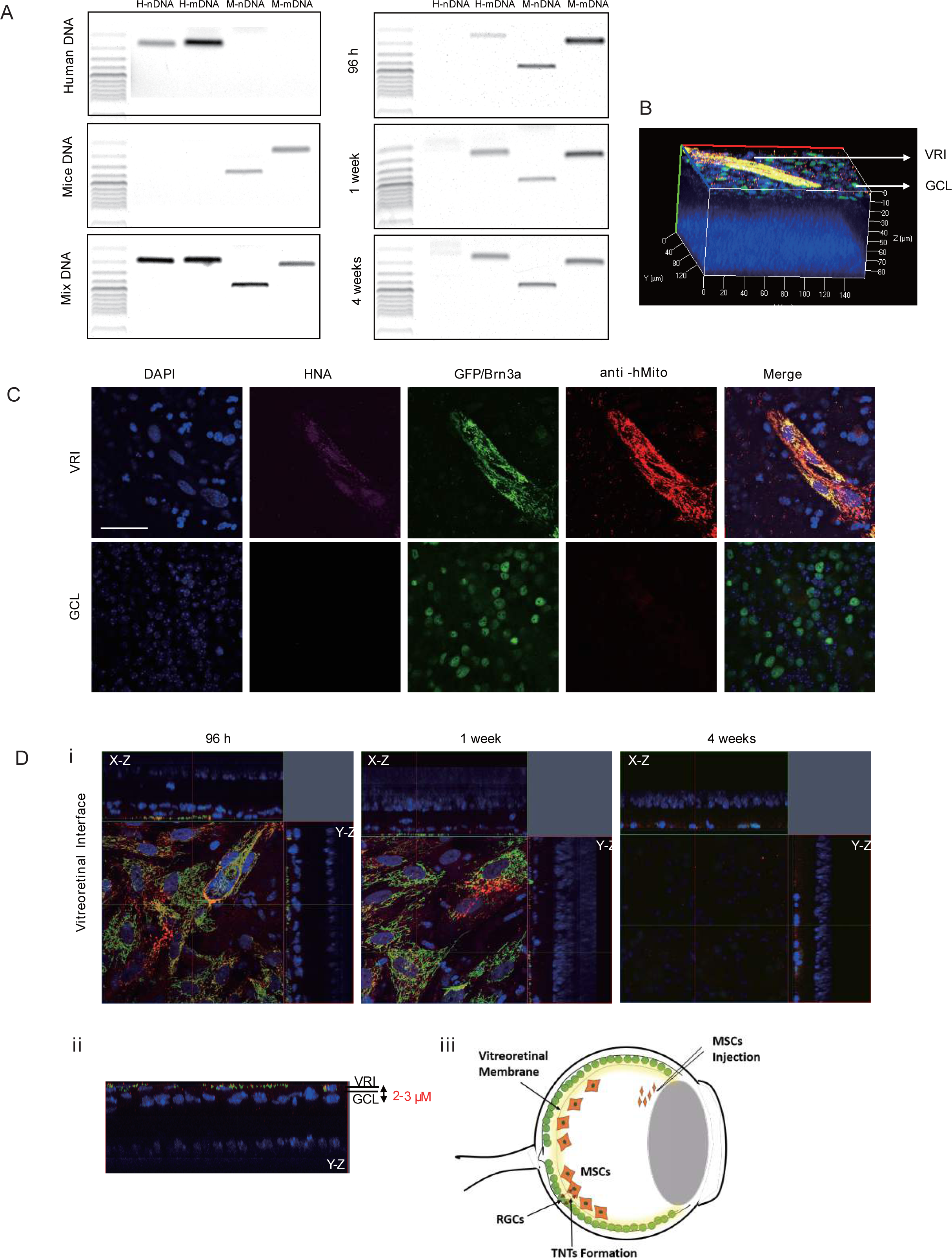
Detection of human mitochondrial DNA within mice retina. **A** PCR results of retina tissue at 96 h, 1 week and 4 weeks post treatment. DNA from human iPSC-MSCs, mice tissue and mixture (human cells + mice tissue) served as control. (H-nDNA: human specific nuclear DNA, H-mDNA: human specific mitochondrial DNA, M-nDNA: mice specific nuclear DNA, M-mDNA: mice specific mitochondrial DNA). **B** Z-stack reconstructed images show that MSC cells were detected on Vitreoretinal Interface (VRI). **C** Retina were stained with anti-GFP (Green), RGC marker Brn3a (Green), anti-human mitochondrial marker anti-hMito (Red), human nuclear maker HNA (Purple) and DAPI (Blue). 1 week post iPSC-MSC injection, Vitreoretinal Interface (VRI) and retinal ganglion cell layer (GCL) were analyzed. **D** i) Z-stack images show duration of stay of iPSC-MSCs on vitreoretinal interface. ii) Reconstructed Y-Z image shows the location of iPSC-MSCs 1 week after injection. iii) A model shows the injected iPSC-MSCs form TNTs and transfer mitochondria to RGCs, passing through the vitreous retina barrier.

To clarify how iPSC-MSCs donate mitochondria to RGCs and penetrate the inner limiting memberane, we tracked iPSC-MSCs located in the eyeball. We established that iPSC-MSCs could be detected on the surface of the vitreoretinal membrane 96 hours and 1 week after injection (Figure 3D). These intravitreal injected iPSC-MSCs were tightly attached on the surface of the vitreoretinal membrane and formed almost a single layer (Figure 3Di). Z-stack images revealed that the layer of RGCs was 2-3 μM beneath the iPSC-MSCs attached on the inner limiting membrane (Figure 3Dii). It is possible that MSC-formed TNTs with diameter of 30-200 nm and length of 30 μM could penetrate the barrier and connect with RGCs for mitochondria transfer (Figure 3Diii). Another possibility is that the astrocyte and the end foot of Muller cell in the limiting membrane transported mitochondria into the RGC layer. Although mitochondria from iPSC-MSCs remained detectable until 4 weeks after injection, we failed to detect any iPSC-MSCs on the surface of the vitreoretinal membrane (Figure 3D). This concurs with previous reports of short-term survival of iPSC-MSCs following intravitreal transplantation (Hill et al., 2009; Labrador Velandia et al., 2018).

### Prevention of RGC loss with iPSC-MSCs treatment in Ndufs4 deficient mice

To determine whether iPSC-MSC administration prevents RGC loss in an Ndufs4 mouse model, we performed intravitreal injection of iPSC-MSCs into 3 week old Ndufs4 KO mice. Age and sex matched KO mice were divided into three groups randomly and subjected to one of the following: iPSC-MSC treatment (KO+MSC), vehicle treatment (KO+PBS), iPSC-MSC pre-treated with rotenone to induce dysfunctional mitochondria (KO+MSC(R)). WT littermate mice with vehicle administration served as controls (WT+PBS). After 1 week of treatment (4 week old mice), number of RGCs did not differ significantly between WT+PBS, KO+PBS and KO+MSC group, confirming that RGC loss could not be detected until week 6 (Figure 1A). Surprisingly, compared with the KO+MSCs group, RGCs in the KO+MSC(R) group were significantly reduced (Figure 4A, p<0.05) even less than the control group. These results provide an important warning that transplantation of mitochondrial dysfunctional MSCs no longer has the capacity for RGC protection and are also harmful to RGC homeostasis, leading to RGC loss. By 4 weeks after iPSC-MSCs injection (mice at 7 weeks of age), whole retina analysis revealed that Ndufs4 KO mice with vehicle administration (PBS group) presented a significant reduction in RGCs compared with WT mice with PBS treatment (Figure 4A,p<0.001). In contrast, iPSC-MSC administration significantly prevented RGC loss compared with vehicle treatment in Ndufs4 KO mice (Figure 4A, p<0.05). Of note, compared with their WT counterparts, MSC(R) treatment failed to protect RGCs from death in KO mice and only around 1/2 of all RGCs remained by 4 weeks post transplantation.

**Figure 4.**
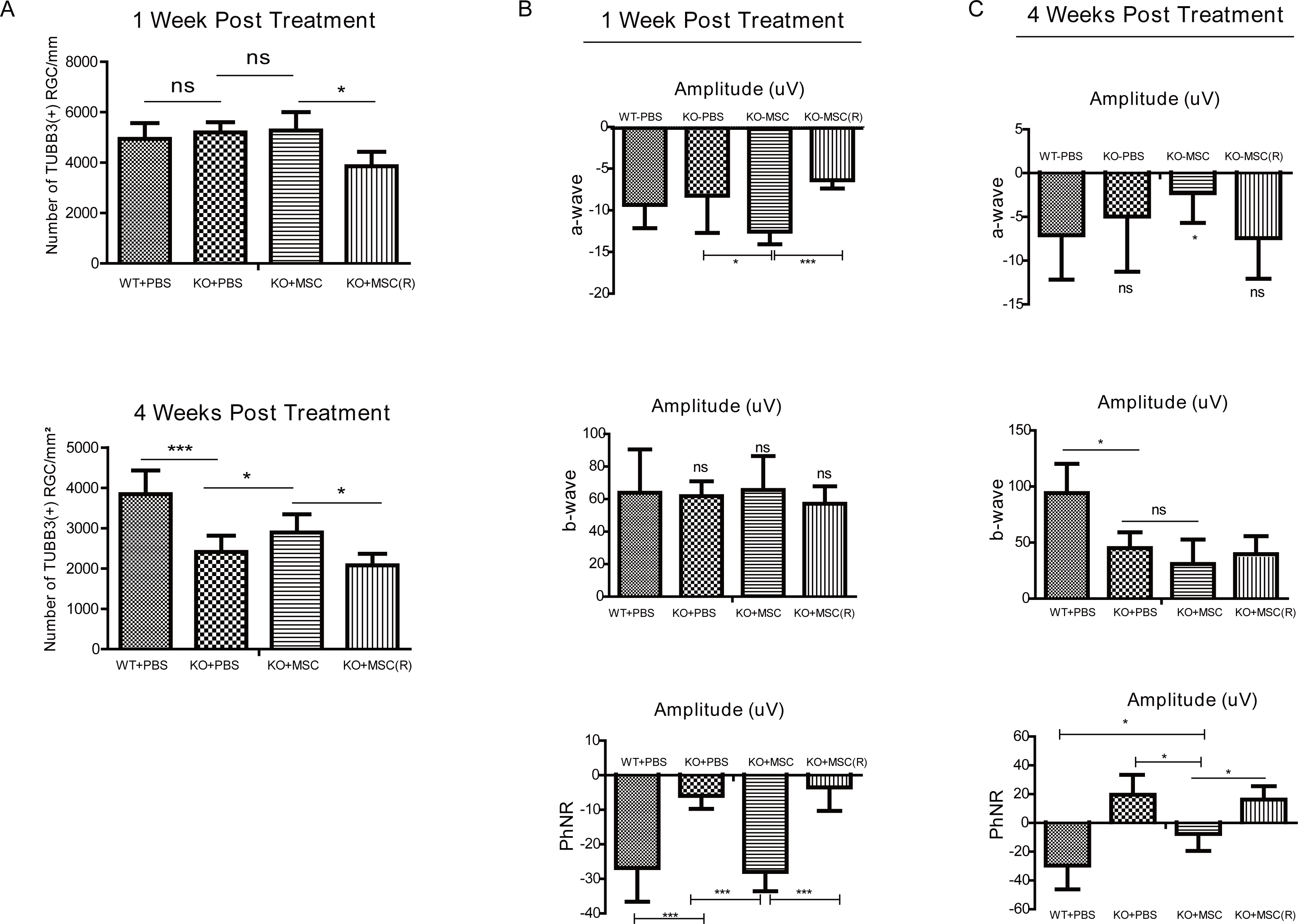
iPSC-MSCs administration prevents retinal ganglion cell and functional loss. **A** Density of RGC in the RGCL immune labeled with TUBB3 antibody in wild type mice and KO mice treated with PBS, MSC and MSC(R) were compared both at one week (upper lane) and four weeks (lower lane) after intravitreal injection. (n=5). B-C Retinal electrography of the mice at one week (B) and four weeks (C) after intervitreal treatment. The amplitude of a-waves (upper lane), b-wave (middle lane) and PhNR (lower lane) was quantified (n=10). Data information: In (A–C), data are presented as mean ± SD. *p≤0.5, **p≤0.01, ***p≤0.001, ns, not significant (One-way ANOVA,).

### Administration of iPSC-MSCs prevents functional decline of RGCs in an Ndufs4 mouse model

In *Ndufs4* knockout mice, ERG measurement showed significant functional impairment of the retina prior to the development of neurodegeneration with detectable cell loss (Figure 1B). We therefore determined whether iPSC-MSC-based treatment could improve the electrical response of the mouse retina with defective mitochondria. Waveforms derived from photoreceptors (a-wave), the inner retina (b-wave) and RGCs (PhNR) were recorded.

At 1 week post treatment, the KO-MSC and KO-PBS group showed no significant differences in the changes to a-wave and b-wave amplitudes compared with age-matched WT mice (WT-PBS) (Figure 4B). Nonetheless compared with the KO-PBS group, KO-MSC mice showed significantly increased a-wave amplitudes (Figure 4B). Moreover, the PhNR amplitude in KO-PBS mice showed a significant decrease compared with that of WT-PBS control mice (Figure 4B, p<0.001), implying that malfunctions of RGCs occurred prior to cell death. Importantly, iPSC-MSC treatment effectively preserved the amplitude of PhNR in KO mice although such preservation was largely diminished when iPSC-MSCs pre-treated with Rotenone were administered (Figure 4B, p<0.001 vs KO-MSC group), indicating that iPSC-MSCs with dysfunctional mitochondria were no longer protective for RGCs. In addition, intravitreal injection of iPSC-MSCs or PBS had little effect on the peak of a-wave, b-wave and PhNR (Data not shown). These data imply that iPSC-MSC treatment at week 1 attenuated the decrease in PhNR amplitudes and increased the a-wave amplitude in Ndufs4 mice with mitochondrial defect.

At week 4 post treatment, there was a significant difference in b-wave and PhNR-wave amplitudes between WT-PBS mice and KO-PBS mice (Figure 4C). Compared with WT-PBS mice, KO-PBS mice displayed more severely diminished b-wave and PhNR-wave amplitudes. In addition, KO-PBS mice showed a prolonged time to the peak of b wave. These data provide evidence that retinal degeneration progressively occurred in KO mice. In contrast, iPSC-MSC treatment effectively prevented this progressive decrease in PhNR amplitude of KO mice (Figure 4C). Nonetheless when iPSC-MSCs were pre-treated with Rotenone (KO-MSC(R)), no protective effect could be detected in PhNR amplitudes. This indicates that only healthy mitochondria transferred from donor cells could improve the function of RGCs. Taken together, RGCs (PhNR) benefit from transferred healthy mitochondria until 4 weeks following iPSC-MSC administration.

### Müller cell activation and inflammatory response in whole retina of Ndufs4-diffecient mice

It has been reported that glial fibrillary acidic protein (GFAP) is normally expressed in astrocytes in WT mice. In response to retinal injury, GFAP gene transcription is strongly activated in the müller cell. To evaluate müller cell activation following iPSC-MSC based therapy in complex I defect mice immunohistochemistry of GFAP was performed. GFAP positive cells were limited in Müller cell end-feet and astrocytes in the nerve fiber layer and ganglion cell layer (GCL) of the retina in WT control mice (Figure 5A). By week 7, the GFAP positive signal in KO mice had shown an axial extent throughout the retina with an increased intensity (Figure 5A-B). As shown in figure 5A-B, 4 weeks post iPSC-MSC treatment, fluorescent intensity of GFAP staining had obviously declined in KO mice retina compared with untreated KO mice. These data indicate that iPSC-MSCs could reduce abnormal activation of Müller cells in the mouse retina with defective mitochondria.

**Figure 5.**
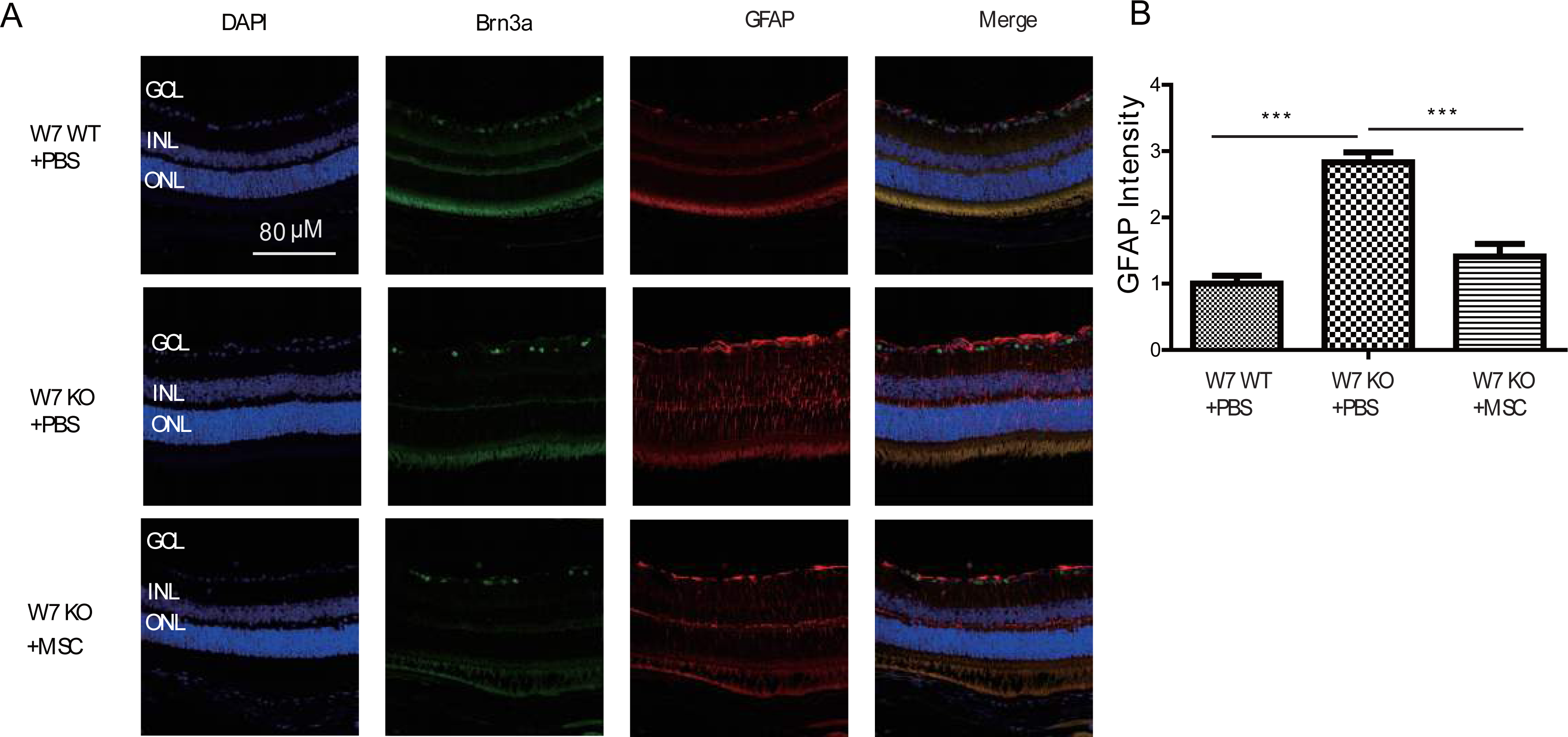
Muller cell/astrocyte activation under iPSC-MSC transplantation. **A** Retinal sections from WT, KO-PBS-injected, and KO-MSC-injected eyes were stained with Brn3a (green) and GFAP (red) antibodies. Sections were counterstained with DAPI (blue). Scale bars: 80 μM. (GCL: ganglion cell layer; INL: inner nuclear layer; ONL: outer nuclear layer). **B** GFAP intensity was analyzed. (n=10, Error bar represent SD, p-values calculated using One-way ANOVA, *** p<0.001).

Results of Cytokine protein arrays and hierarchical clustering analysis of differential protein expression unveiled that compared with week4 WT mice (W4WT+PBS), KO mice (W4KO+PBS) are biased to an inflammatory status as evidenced by remarkably increased pro-inflammatory cytokines (i.e. CD30L, IL-5, TNFα, IFNg etc.) but decreased anti-inflammatory cytokines and cell survival factors (i.e. IL-10, ICAM1etc.)(Figure EV5 & Table EV1). Notably, many of these prominently increased cytokines such as IL-17, IL-1b TNFα, and CD30L (Abdalla et al., 2004; Bonizzi and Karin, 2004; Franchi et al., 2006; Infante-Duarte et al., 2000; Jovanovic et al., 1998; Yue et al., 2005) are associated with activation of NF-κB signaling pathway, a master signaling pathway in the regulation of inflammation. Importantly, one week post iPSC-MSC treatment, the enhanced pro-inflammatory cytokines had remarkably declined in the retina of KO mice (W4KO+MSC). Nonetheless rotenone-treated iPSC-MSCs (W4KO+MSC(R)) lost protective ability when compared with the W4KO-MSC group. Of note, pro-inflammatory cytokines of W4KO-MSC(R) mice were much higher than those of W4KO-PBS mice, indicating that MSCs with dysfunctional mitochondria have lost anti-inflammatory properties instead are harmful to the retina (Figure EV5 & Table EV1).

We carried out four comparisons, KO+PBS v.s. WT+PBS, KO+MSC v.s. KO+PBS, KO+MSC(R) v.s. KO+PBS, and KO+MSC v.s. KO+MSC(R) under two time points. A total of 22 proteins that display 1.50 or above folds of difference in at least one of these comparisons were identified. By hierarchical clustering analysis on expression of these differential proteins using Cluster 3.0, we detected three clusters of cytokines and treatment groups with similar profile (Figure 6). As shown at Figure 6, KO mice with PBS or MSC(R) treatment at week 4 shared majority of commonly increased pro-inflammatory cytokines, such as IL-5, IL-17, GM-CGF, IL-6 etc. indicating that increased inflammation in KO mice cannot be suppressed by MSC(R) treatment. By contrast, week 4 mice subjected to KO-MSC treatment (panels 3, 4) largely reversed these increased pro-inflammatory cytokines, supporting effective protection of MSC treatment in reduction of excessive inflammation. Similar dynamics of inflammation was observed between PBS and MSC(R) treatment at week 7 KO mice (panels 5, 6), further supporting inefficacy of MSC(R) treatment. However, the protective effects of MSC, but not MSC(R) were still evidenced up to week 7 (panels 7, 8). This result indicates distinct inflammatory status of WT and KO mice, and different therapeutic efficacy of MSC and MSC(R).

**Figure 6.**
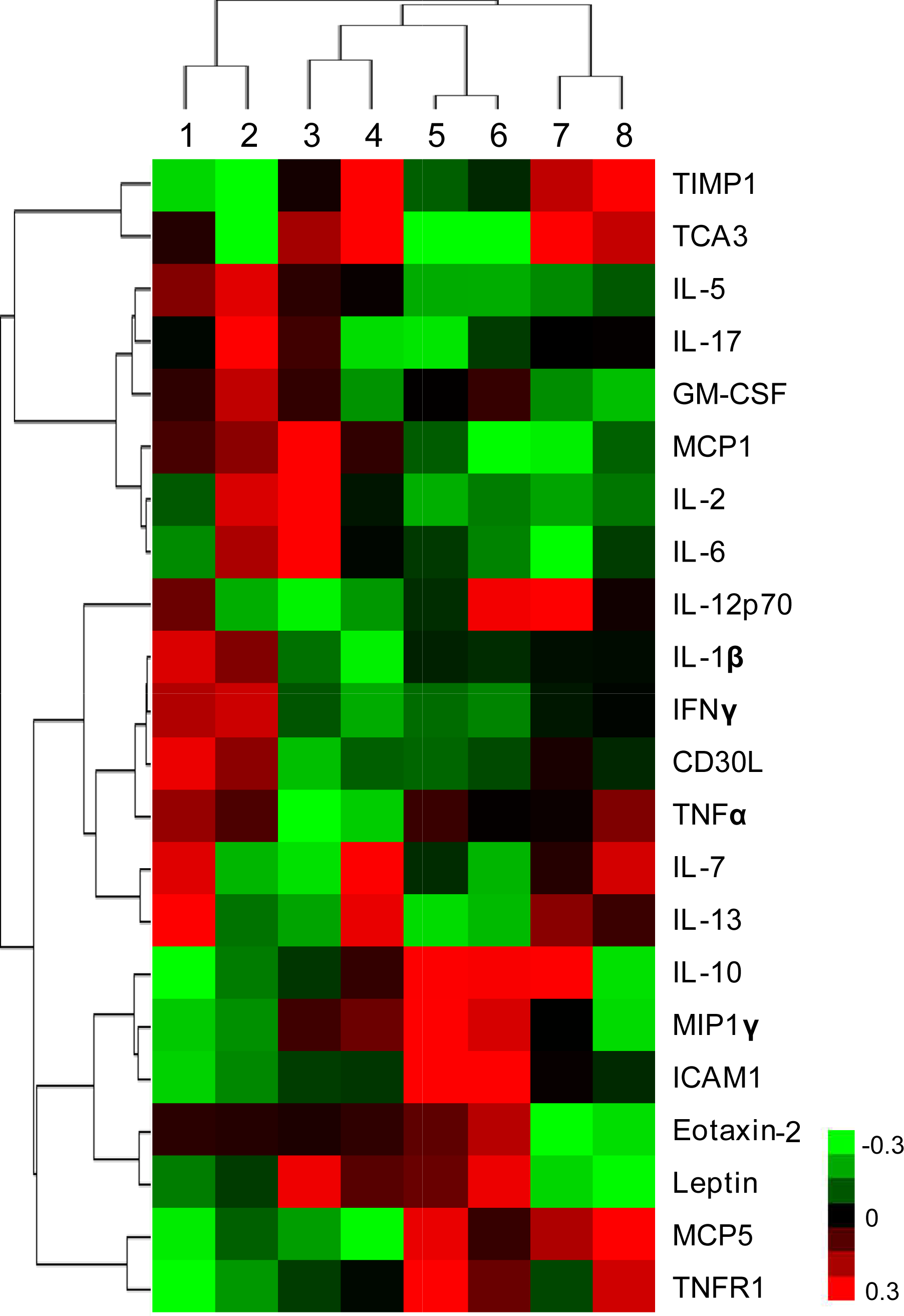
Cytokine protein arrays and hierarchical clustering analysis of Ndufs4 Mice. Hierarchical clustering of 22 differential proteins having at least 1.50 fold difference from one of the four comparisons, KO+PBS v.s. WT+PBS (panels 1), KO+MSC v.s. KO+PBS, KO+MSCr v.s. KO+PBSs, and KO+MSC v.s. KO+MSCr at week 4 (W4) and week 7 (W7). Over- and under-expressed proteins are indicated by red and green, respectively. Panels 1: W4-KO+PBS/WT+PBS; Panels 2: W4-KO+MSCr/KO+PBS; Panels 3: W4-KO+MSC/KO+PBS; Panels 4: W4-KO+MSC/KO+MSCr; Panels 5: W7-KO+PBS/WT+PBS; Panels 6: W7-KO+MSCr/KO+PBS; Panels 7: W7-KO+MSC/KO+PBS; Panels 8: W7-KO+MSC/KO+MSCr.

## Discussion

In this study, we have demonstrated that retinal ganglion cells (RGCs) display functional defects and eventually death in a *Ndufs4* knockout mouse model; iPSC-MSCs delivered by intravitreal injection successfully donated mitochondria to RGCs; iPSC-MSC-mediated mitochondrial transfer could prevent RGC loss, preserve RGC function, and help to manage abnormal activation of Müller Cells and inflammatory state of the whole retina.

iPSC-MSCs could transfer their healthy mitochondria to the retina with a hereditary mitochondrial disorder via intravitreal injection. Additionally, the mitochondrial transfer was detected 96 hours following injection. The vitreous chamber has been shown to be a favorable microenvironment for iPSC-MSC survival; iPSC-MSCs remained in the vitreous cavity and had not integrated into the retina 4 weeks post treatment. PCR results confirmed that only mitochondria of iPSC-MSCs could be transferred into the retina of mice after administration and none fused into the retina of Ndufs4 mice. Our results are consistent with the data of Johnson et. al (Johnson et al., 2010b) who observed extremely poor retinal integration of MSCs following intravitreal injection. In some studies, the authors showed that injected cells remained in the vitreous cavity and did not integrate into the retina 12 weeks after treatment (Ezquer et al., 2016). Due to the limited lifecycle of this mouse model (around 7 to 9 weeks), we could not evaluate the maximal time for mitochondria transfer from iPSC-MSCs maintained within mice retina.

The donated mitochondria were mainly located in the RGC layer. Retinal ganglion cells are responsible for propagating signals derived from visual stimuli in the eye to the brain, along their axons within the optic nerve to the superior colliculus, lateral geniculate nucleus and visual cortex(Mead and Scheven, 2015). Although only a few mitochondria were detected within the inner nuclear layer (INL) and inner plexiform layer (IPL), it is worth investigating whether donated mitochondria can move along the axon of optic nerves from RGCs to the brain and protect against neurodegeneration in the central neural system. It is also worth to study the possibility of MSC connected with astrocyte or Muller cell and transfer mitochondrial to them.

The mechanisms by which iPSC-MSCs support retinal regeneration are thought to be mainly via paracrine effects (Cejkova et al., 2013; Kureshi et al., 2015). Our study indicates that when iPSC-MSCs were pretreated with complex I inhibitor (Rot), RGC loss could not be prevented. It suggests that only healthy mitochondrion donated from MSCs directly protect RGC against hereditary mitochondrial damage. Hence, mitochondrial transfer, as another important therapeutic mechanism, plays an important role in iPSC-MSC-based treatment. In addition, injured MSCs or those with aging mitochondria may not be suitable as therapeutic donor cells.

The remarkable ability of iPSC-MSCs to prevent RGC loss after intravitreal transplantation has been corroborated by ERG study, one of the most accepted techniques to evaluate retinal electrical responses (Ezquer et al., 2016). Our results imply that the PhNR derives from retinal ganglion cells that were protected when treated with iPSC-MSCs both 1 and 4 weeks post injection. In addition, 1 week after treatment, the amplitude of PhNR was increased in KO mice, although the number of RGS was unchanged. This suggests that functional improvement of RGCs occurs before an increase in cell number following iPSC-MSC treatment.

Mitochondrial complex I dysfunction in the retina is strongly related to an innate immune and inflammatory response that results in loss of retinal ganglion cell function and death, as in Leber’s hereditary Optic Neuropathy (Yu et al., 2015). Four weeks following iPSC-MSC treatment, glial fibrillary acidic protein (GFAP) expression obviously declined in the retina of KO mice. These results indicated that iPSC-MSCs could reduce abnormal activation of glial cells in the mouse retina with defective mitochondria. High levels of GFAP have been associated with pathological inflammation and RGC loss (Kurihara et al., 2006). Alfred et al. reported that increased innate immunity, microglial and astrocyte transcripts as the activating factors occurred before functional defects as well as cell loss in Ndufs4 mice (Yu et al., 2015). Cytokines protein-array revealed that proinflammatory cytokines were clearly increased in the retina of KO mice at week 4 compared with WT. It has been suggested that activation of microglia with IFN-g, GM-CSF and MCP-1 leads to neuronal injury (Lin and Levison, 2009; Meda et al., 1995; Vargas et al., 2005). In turn, activated microglia produced excessive IL-1b and TNF α that are toxic to neurons (Block and Hong, 2005). These results demonstrate a link between mitochondrial dysfunction (*Ndufs4* mutation) and inflammation that contributes to neural degeneration. No doubt, management of the inflammatory response will be critical to RGC survival with complex I defect.

iPSC-MSC treatment in *Ndufs4*-KO mice effectively reduced neuro-destructive cytokines such as TNF α, MIP−1g, GM-CSF, IL-5, IL-17 and IL-1b (Murray and Wynn, 2011; Yuan and Neufeld, 2000). In addition, it has been reported that TNF α stimulates the formation of TNTs and promotes mitochondrial transfer (Zhang et al., 2016). There is evidence of an obligatory role of NF-κB signaling in the regulation of mitochondrial transfer via TNT formation (Ahmad et al., 2014). Interestingly, cytokines that were increased in Ndufs4-KO mice and down-regulated by iPSC-MSC treatment, such as GM-CSF, MCP-1, IL-17, IL-1b TNF-α, IL−12p70 and CD30L, were related to NF-κB pathway (Abdalla et al., 2004; Bonizzi and Karin, 2004; Franchi et al., 2006; Infante-Duarte et al., 2000; Jovanovic et al., 1998; Yue et al., 2005). High production of cytokines such as TNF α and activation of NF-κB signaling in KO mice might be the trigger for mitochondrial transfer. We found that Rotenone pretreated MSC lost their protective effect, indicating that only the transferred healthy mitochondria were of benefit to the retina, besides the paracrine effects. This is consistent with previous study using Transwell units that the crosstalk between paracrine action and mitochondrial transfer is an interaction of two independent processes with consequent MSC-mediated cell protection (Vallabhaneni et al., 2012).

In summary, **r**etinal mitochondrial defects provoke pathological inflammation, RGC dysfunction and cell death. Intravitreal transplantation of iPSC-MSCs can effectively donate functional mitochondria to RGCs, overcoming the barrier of the inner limiting membrane of the retina. The donated mitochondria of iPSC-MSCs can rescue RGC function against mitochondrial damage-induced excessive inflammation and retinal degeneration (Figure 7). We believe intravitreal transplanted MSCs can donate functional mitochondria to RGCs. Application of stem cell mitochondrial transfer opens new possibilities to prevent or halt RGC degeneration responsive to acquired or hereditary mitochondrial retinal disorders.

**Figure 7.**
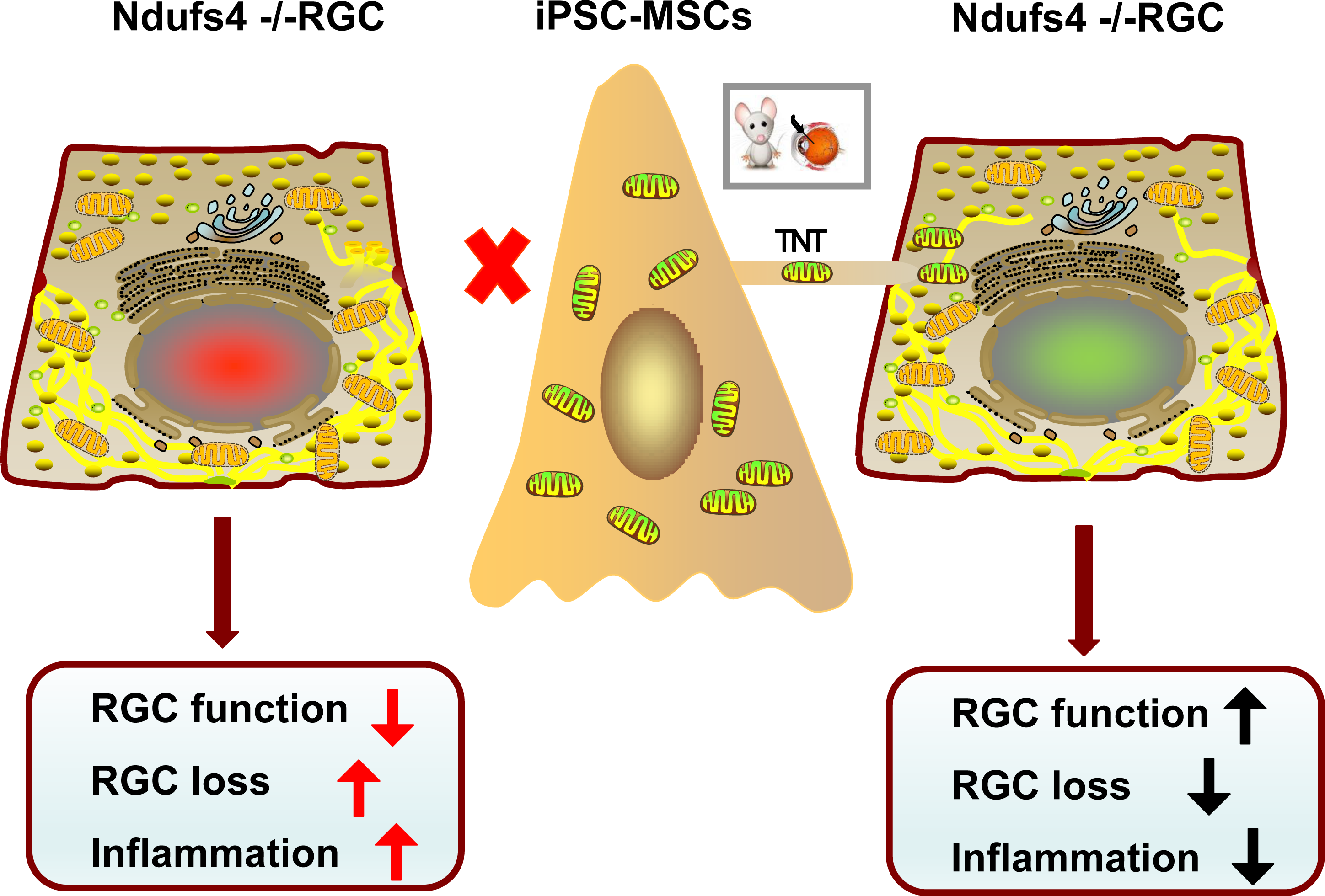
The schematic cartoon indicating the mitochondrial damage-provoked RGC degeneration and the roles of iPSC-MSCs treatment in RGC protection.

## Methods

### *In vitro* cell culture and labeling

Human iPSC-MSCs (Lian et al., 2010) as the mitochondria donor cells were prepared and cultured in Dulbecco modified Eagle’s medium (DMEM; Gibco, Invitrogen, Carlsbad, CA, USA) plus 15% fetal bovine serum (FBS), 2 ng/ml EGF and 2 ng/ml FGF (Lian et al., 2010). They were then transfected with Lentiviral-Mitochondrial-Green Fluorescence Protein (LV-Mito-GFP) (Jiang et al., 2016; Zhang et al., 2016). iPSC-MSCs pretreated with mitochondrial complex I inhibitor Rotenone (R) named as iPSC-MSCs(R) and served as the mitochondria viability control group (Jiang et al., 2016).

Primary cultures of RGCs were prepared from the isolated retina (*n* ≥ 24/preparation) of 2 to 4 day old mice (Sappington et al., 2006). CD90.1 MicroBeads (Miltenyi Biotec) were used to isolate RGCs from cell suspension. For the co-culture experiment with iPS-MSCs, RGCs as the recipient cells were labeled with CellTrace Violet (Excitation:405, C34557, Invitrogen)

### Ndufs4 Knockout Mouse Model

All experiments on mice were approved by the committee on the Use of Live Animals in Teaching and Research (CULATR) of the University of Hong Kong. Mice were housed in a Minimal Disease Area (MDA) under controlled environmental conditions on a 12 □ hrs light-dark cycle with *ad libitum* access to food and water at room temperature, 22°C. C57BL/6 mice were crossed with *Ndufs4+/-* mice for colony maintenance. To acquire *Ndufs4-/-* (KO) and *Ndufs4+/+* (WT) mice, heterozygous (*Ndufs4+/-*) male mice were mated with female mice.

### Intravitreal Injection

To determine whether iPSC-MSCs could exert their neuroprotective effect in Ndufs4-/- mice via intercellular mitochondrial transfer, iPSC-MSC treatment was performed via intravitreal injection. iPSC-MSCs or PBS and iPSC-MSCs(R) were injected into the vitreous body of mice at age 3 weeks. A cell suspension containing 1□ × □ 10^4^ iPSC-MSCs/ iPSC-MSCs(R) in 0.5 μL PBS was slowly injected into the vitreous cavity. The same volume of PBS was injected as the placebo control group. At 96 hrs, 1 week and 4 weeks after injection, mice were sacrificed for examination. Eyes with either cataract or vitreous hemorrhage after the injection were excluded from the study.

### Electroretinography (ERG)

To evaluate the visual function of WT and KO Ndufs4 mice, scotopic flash electroretinography (ERG) was performed. To demonstrate the effect of iPSC-MSC treatment, ERG was performed 1 and 4 weeks after injection. After overnight dark adaption, mice were anesthetized by 1:1 mixture of ketamine and xylazine, pupils were dilated using mydriasis. Both eyes were anesthetized topically with 0.5% proparacaine (Alcon, US), ring shape gold electrodes were placed closely contact the cornea of both eyes and a ground electrode was placed subcutaneously by the tail. After 10 min adaption, eyes were exposed to a series of increasing intensity light flashes. Stimulus intensities of 11.38 scd/m^2^, 16.60 scd/m^2^ and 22.76 scd/m^2^ were applied on a green background to record the response of the eye. Results were analyzed for a-wave, b-wave, and PhNR by ERG View v4.380R software.

### Immunofluorescence labeling of RGC in Whole Mount Retina

To evaluate RGC loss, RGC labeled with POU domain protein 3A (Brn3a) or Tubulin beta 3 (TUBB3) were counted in whole mount retina. At different time points, eyes were enucleated and fixed in 4% paraformaldehyde in PBS for 30 mins at 4 °C followed by a cornea puncture and extra 30 mins fixation. Next, the anterior segment was removed and the retina detached carefully from the underlying RPE and sclera. After soaking with 5% BSA and 0.5% Triton-X-100 in PBS for 1 hrs, the retinas were incubated with a 1:500 dilution of first antibody at 4 °C overnight and a 1:1000 dilution of secondary antibody at room temperature for 2 hours (Brn3a (Santa Cruz Biotechnology, TX, USA), TUBB3 (Biolegend, CA, USA)). After washing, retinal tissue was mounted with mounting medium. Sixteen individual fluorescent images from four regions per retina were taken using the fluorescent microscope: central retinal (C-R), near-peripheral retina (N-P), mid-peripheral retina (M-P) and far-peripheral (F-P) retina. (Leica laser microscope 700).

### Morphometric analysis of the vertical retinal sections

Eyes from KO and WT mice at 3, 4 and 7 weeks old were collected and processed for paraffin vertical sections, followed by standardized Hematoxylin and Eosin (H&E) staining protocol. H&E retinal images were taken from 2 retinal areas: Central retina (C) and peripheral retina (P). The thickness of inner nuclear layer (INL), inner plexiform layer (IPL) and outer nuclear layer (ONL) were measured by ImageJ software.

### Assessment of mitochondrial transfer and Müller Cell Activation

Mitochondrial transfer and Müller Cell Activation were detected via immunofluorescence on flat mounted retina. To observe mitochondrial transfer within the retina following iPSC-MSC injection, TUBB3/Brn3a was used to locate RGCs and anti-GFP/anti-hMito were used to label mitochondria from iPSC-MSCs (Anti-hMitochondria (Abcam, CA, USA), Anti-GFP (Invitrogen, CA, USA), human nuclear antigen (HNA; Millipore)). To evaluate Müller Cell activation following iPSC-MSC based therapy, immunohistochemistry of GFAP (Sigma, MO, USA) was performed. Samples were examined using a laser scanning confocal microscope LSM700 or a time-lapse video recorder.

### Polymerase chain reaction (PCR)

Total DNA of the retina was extracted using a DNA kit (Qiagen, 74104). 1 µg of total DNA was used as a template for PCR. PCR cycles were as follows: 94 °C for 5 mins, 35 cycles at 94 °C for 20 seconds, 57 °C for 30 seconds, 72 °C for 40 seconds, and a final extension at 72 °C for 5 mins. The amplified products were analyzed by electrophoresis in 1.0% agarose gel containing ethidium bromide. The primers used in this study were Human specific mtDNA, Forward 5’-ACCACCCAAGTATTGACTCACC-3’ Reverse 5’-GGGGGTTTTGATGTGGATTGG-3’; Murine specific mtDNA, Forward 5’-AGGCATGAAAGGACAGCACA-3’ Reverse 5’-TTGGGGTTTGGCATTAAGAGGA-3’; Human specific nDNA, Forward 5’-GCATCGTGCTGCCTTGATTT-3’ Reverse 5’-CCGTAAGAGGCCAGCACATT3’; Murine specific nDNA, Forward 5’-GAATTCAGATTTGTGCATACACAGTGACT-3’and Reverse 5’-AACATTTTTCGGGGAATAAAAGTTGAGT-3’.

### Quantified inflammatory cytokines array of retina

Retinas were isolated from mice and snap frozen for analysis of inflammatory cytokines. The concentration of 40 cytokines was determined using the Quantibody Mouse Inflammation Array Kit according to the manufacturer’s instructions (RayBiotech, Inc., Norcross, GA, USA). The array was designed to quantitatively detect 40 inflammatory cytokines: BLC (CXCL13), CD30 Ligand (TNFSF8), Eotaxin-1 (CCL11), Eotaxin-2 (MPIF-2/CCL24), Fas Ligand (TNFSF6), GCSF, GM-CSF, I-309 (TCA-3/CCL1), ICAM-1 (CD54), IFN-gamma, IL-1 alpha (IL-1 F1), IL-1 beta (IL-1 F2), IL-10, IL-12 p70, IL-13, IL-15, IL-17A, IL-2, IL-21, IL-3, IL-4, IL-5, IL-6, IL-7, KC (CXCL1), Leptin, LIX, MCP-1 (CCL2), MCP-5, M-CSF, MIG (CXCL9), MIP-1 alpha (CCL3), MIP-1 gamma, Platelet Factor 4 (CXCL4), RANTES (CCL5), TARC (CCL17), TIMP-1, TNF alpha, TNF RI (TNFRSF1A) and TNF RII (TNFRSF1B). Signals (green fluorescence, Cy3 channel, 555 nm excitation, and 565 nm emission) were captured using a GenePix 4000B laser scanner (Bio-Rad Laboratories, Hercules, CA) and extraction performed with GenePix Pro 6.0 microarray analysis software. Quantitative data analysis was performed using RayBiotech mouse Inflammation Array 1 software (QAM-INF-1_Q-Analyzer). Sample concentrations (pg/ml) were determined from mean fluorescence intensities (median values) compared with five-parameter linear regression standard curves generated from standards provided by the manufacturer.

### Statistical analysis

Statistical analysis was performed using Prism 5.04 Software (GraphPad Software for Windows, San Diego, CA, USA), and results are reported as mean ± S.D. Comparisons between more than two groups were analyzed by one-way ANOVA test. Comparisons between two groups were performed using Student’s t-test. A p-value <0.05 was considered significant.

## Acknowledgement

This research was in part supported by National Natural Science Grant of China (No 31571407, 31270967 to Q Lian); Hong Kong Research Grant Council General Research Fund HKU17113816, HKU772510M to Q Lian); and the key grant from the Science and Technology Foundation of Guangdong Province of China (2015B020225001)

## Author Contributions

D J: experiments, collection and/or assembly of data and data analysis; .F H, ZZ, LC CM: preparation of MSC and collection of data; CL,SH, YZ collection of data; BY, P C H Tse data analysis; Q F, K C data analysis and manuscript writing; Q L: concept and design, dana analysis, manuscript writing, and final approval of the manuscript.

## Competing Interests

The authors have declared that no competing interest exists.

## The Paper Explained

### Problem

Retinal ganglion cell (RGC) degeneration is extremely difficult to repair and is often associated with mitochondrial dysfunction. Transplantation of mesenchymal stem cell (MSC) holds promise to protect RGC but beneficial mechanisms are largely unknown.

### Results

Here, we report intravitreal transplantation of iPSC-MSCs could overcome the barrier of the inner limiting membrane of the retina, donate functional mitochondria to rescue RGCs from degeneration in mitochondrial *Ndufs4* deficient mice, Importantly; the beneficial effects of mitochondrial donation by MSCs were associated with management of mitochondrial damage-induced excessive inflammation and overall preservation of RGC functions (Figure 7).

### Impact

MSCs modulated-mitochondrial donations effectively protect RGC from degeneration. Application of stem cell mitochondrial transfer opens new possibilities in the treatment of RGC degeneration, enhancing the therapeutic potential for acquired or hereditary mitochondrial retinal disorders(Almasieh et al., 2012; Liu et al., 2010; Ting et al., 2016; Yu-Wai-Man et al., 2011).

## Expanded View Figure legends

**Figure EV1 Immunofluorescent detection of GFAP in KO and WT mice.**

Immunofluorescent images of retina sections with Brn-3a (green), GFAP (red) and DAPI (blue) staining in WT and KO mice at week 3, week4 and week7 postnatal.

**Figure EV2 Strategy of treatment**

**Figure EV3 Coculture iPSC-MSCs and RGCs *in vitro***

iPSC-MSCs were labeled with LV-mito-GFP. RGCs were labeled with CellTrace Violet. Co-cultivation of iPSC-MSCs and RGCs for 24 hrs. Image shown that GFP positive mitochondria were detected within RGCs.

**Figure EV4 iPSC-MSCs administration prevents retinal ganglion cell loss.**

Whole mount retinas were immunolabeled with Brn3a antibody (marker of RGCs) and anti-hMito (marker for mitochondria from human MSC). 96 h, 1week and 4 weeks after treatment, anti-hMito intensity in the ganglion cell layer of the retina were analyzed. (n=5, * p< 0.05)

**Figure EV5 Fold-change (log) of cytokine concentrations in whole retina of Ndufs4 Mice**

A heatmap showing the fold-change (log) of cytokine concentrations in whole retina under different conditions. One row presents one cytokine, one column presents one comparison. The number in figure is the log (fold-change), which is also represented by blue-red color-codes (blue: downregulated; white, no change; red: upregulated). The rows were ordered according to a hierarchical clustering of the fold-change values, by Euclidean distance. Four groups in experiments were compared at two time-points (week 4 and 7 respectively).

Mitochondrial donation by MSCs protects RGC

## Expanded View Table 1 Fold-change of 40 cytokine

Mice at age 3 weeks were treated, 1 week (W4) and 4 weeks (W7) after injection, mice were sacrificed for cytokine examination. Cytokine concentrations of retinas were compared between different treatment conditions presented as fold-changes. The fold changes of cytokine concentrations were listed.

## Expanded View Movie 1

Whole mount retinas were immunolabeled with TUBB3 antibody (marker of RGCs), anti-GFP (marker for mitochondria from human MSC) and DAPI. Z-stack Images were acquired by a laser scanning confocal microscope LSM 700 with a 488□nm argon laser and a 405□nm laser. Z-stack reconstructed 3D movie of retina clearly showed that most of the mitochondria from iPSC-MSCs were located within the ganglion cell layer.

